# A balanced measure shows superior performance of pseudobulk methods over mixed models and pseudoreplication approaches in single-cell RNA-sequencing analysis

**DOI:** 10.1101/2022.02.16.480517

**Authors:** Alan E Murphy, Nathan G Skene

## Abstract

Recently, Zimmerman *et al*.,^1^ highlighted the importance of accounting for the dependence between cells from the same individual when conducting differential expression analysis on single-cell RNA-sequencing data. Their work proved the inadequacy of pseudoreplication approaches for such analysis – This was an important step forward that was conclusively proven by them. A hierarchical single-cell expression simulation approach (<u>hierarchicell</u>) was developed by Zimmerman *et al*.,^1^ to generate non-differentially expressed genes upon which performance was evaluated using the type 1 error rate; the proportion of non-differentially expressed genes indicated as differentially expressed by a model. However, evaluating such models on their type 1 or type 2 error rate in isolation is insufficient to determine their true performance – for example, a method with low type 1 error may have a high type 2 error rate. Moreover, because no seed was set for the pseudo-random number generator used in hierarchicell, the different methods evaluated by Zimmerman *et al*. were done so on different simulated datasets. Here, we corrected these issues, reran the author’s analysis and found pseudobulk methods outperformed mixed models.

**Contact:** Alan Murphy, Nathan Skene

**Code availability:** The modified version of hierarchicell which returns all error metrics, uses the same simulated data across approaches and has checkpointing capabilities (if runs are aborted or crashed) is available at: https://github.com/neurogenomics/hierarchicell.

The benchmarking script along with the results are available at: https://github.com/Al-Murphy/reanalysis_scRNA_seq_benchmark.

## Introduction

Zimmerman *et al*.,^1^ performed a systematic analysis of methods’ type 1 error rates across the 20,000 iterations of 5 to 40 individuals and 50 to 500 cells at a p-value cut-off of 0.05. Plotting the results showed pseudobulk approaches had the lowest type 1 error at every iteration (Supplementary Figure 1). However, as previously outlined, we need to consider both type 1 and type 2 error rate to accurately benchmark the models. Here, we modified Zimmerman *et al*.’s hierachicell approach to simulate both differentially expressed and non-differentially expressed genes. The differentially expressed genes were randomly simulated with a fold change between 1.1 and 10. We further modified hierachicell to correct the seeding of the pseudo-random number generator to enable fair comparisons across models.

## Main

We tested the models using the Matthews Correlation Coefficient (MCC) giving a balanced measure of performance. MCC is a well-known and frequently adopted metric in the machine learning field which offers a more informative and reliable score on binary classification problems^2^. MCC produces scores in [-1,1] and will only assign a high score if a model performs well on both non-differentially and differentially expressed genes. Moreover, MCC scores are proportional to both the size of the differentially and non-differentially expressed genes, so it is robust to imbalanced datasets. We also benchmarked the models using receiver operating characteristics (ROC) curves for different proportions of differentially expressed genes.

Our MCC analysis demonstrates that pseudobulk approaches are the best performing across all number of individuals and cells variations (Figure 1). There is one exception for sum pseudobulk which performs worse than Tobit at 5 individuals and 10 cells. Figure 1 also highlights a trend whereby pseudoreplication models; ‘Modified t’, ‘Tobit’, ‘Two-part hurdle: Default’ and ‘Two-part hurdle: Corrected’ (which take cells as independent replicates) showed degrading performance as the number of cells increase. This is likely due to the over-estimation of power driven by the dependence between cells from the same individual^3^ and agrees with Zimmerman *et al*.’s findings^1^. On the other hand, both pseudobulk approaches; ‘Pseudobulk: Mean’ and ‘Pseudobulk: Sum’, showed improved performance as the number of cells increase. This trend was also noted in two of the other models; ‘GEE1’ and ‘Tweedie: GLMM’.

**Figure 1:**
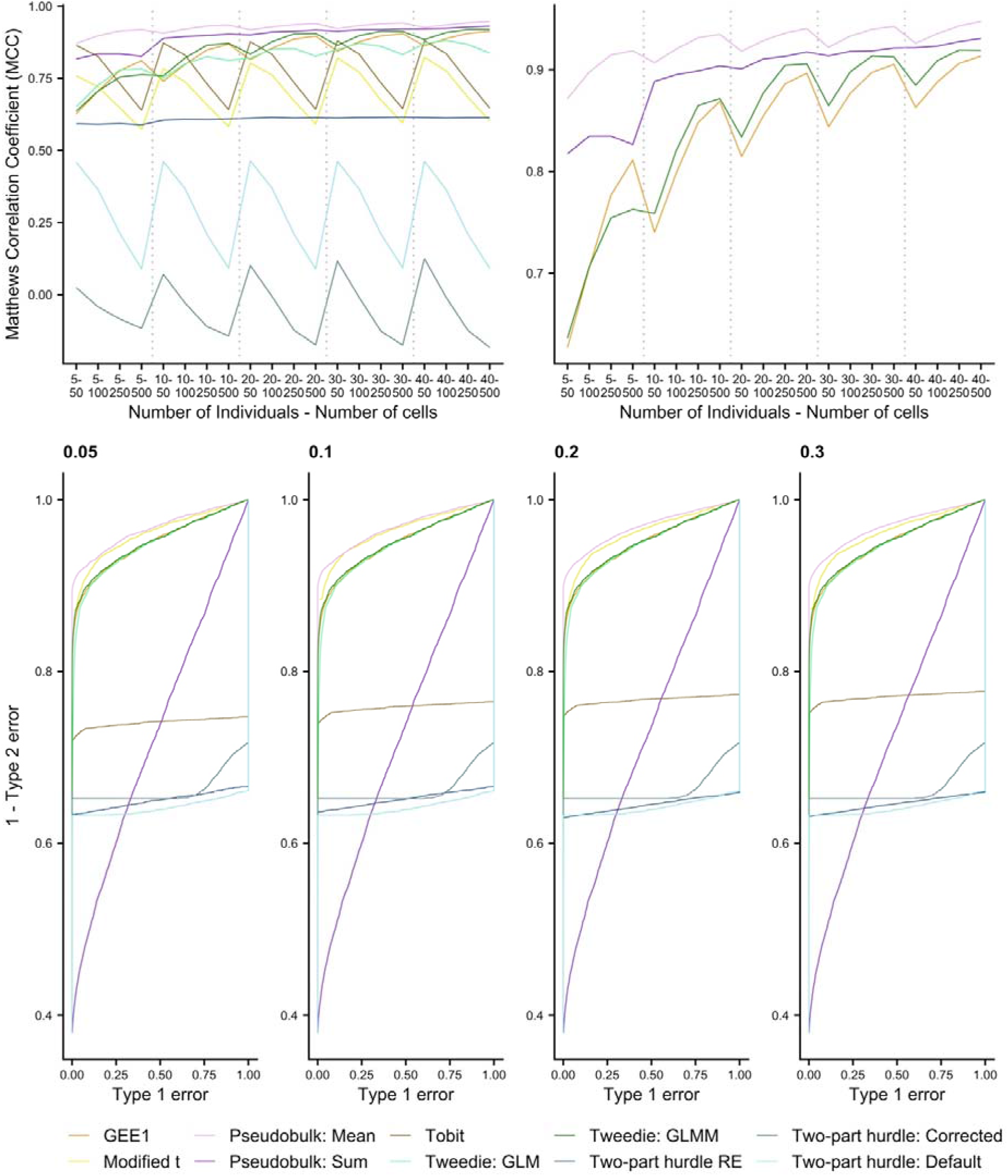
Performance of the analysed models. The top two images give the average Matthews correlation coefficient from the 20,000 iterations; 50 runs for each of the 5 to 40 individuals and 50 to 500 cells at a p-value cut-off of 0.05 on 10,000 genes. Top left shows all benchmarked models whereas right focuses on the top four approaches. The bottom four images give the receiver operating characteristics (ROC) curve across 50 runs each for different proportions of simulated differentially expressed genes (DEGs) - 0.05, 0.1,0.2,0.3. 20 individuals were simulated for case and controls, each with 100 cells. The proportion of DEGs are of the 5,000 non-DEGs simulated. The performance split by each iteration is given in Supplementary Table 2. The different models are pseudoreplication approaches; ‘Modified t’, ‘Tobit’, ‘Two-part hurdle: Default’, ‘Two-part hurdle: Corrected’, ‘GEE1’, ‘Tweedie: GLM’, pseudobulk approaches; ‘Pseudobulk: Mean’, ‘Pseudobulk: Sum’ and mixed model approaches; ‘Tweedie: GLMM’ and ‘Two-part hurdle: RE’. More detail on these models is given in Supplementary Table 1.

Moreover, for statistical test comparisons, another approach is to compare the power of tests at the same test size. That is, to compare the models’ sensitivity (1 – type 2 error) at a consistent type 1 error rate. Therefore, we produced ROC curves for the different approaches, enabling such comparisons. For example, in Supplementary Figure 2, we highlight the different sensitivity scores (1 – type 2 error) of the models obtained at a consistent type 1 error rate of 0.05. Pseudobulk mean was the clear best performing at this type 1 error rate (with a sensitivity >0.9 whereas all other methods had < 0.9) and at all other type 1 error rates (Figure 1). Interestingly, the two mixed model approaches (‘Two-part hurdle: RE’ and ‘GLMM Tweedie’) performed relatively poorly even compared to some pseudoreplication approaches. This analysis demonstrated how pseudobulk mean obtained low type-2 error rates even at the lowest type-1 error rates of the methods benchmarked, supporting our MCC results.

Zimmerman *et al*. argued that pseudobulk methods are “overly conservative” relative to mixed models in their work. The authors here refer to pseudobulk approaches’ lower than nominal levels of type 1 error rates. This was demonstrated in their results where, based on a consistent p-value cut-off of 0.05, they benchmark methods’ performance at identifying non-differentially expressed genes (Supplementary Figure 1). Their analysis showed pseudobulk approaches’ type 1 error rates were below the expected 0.05 of false positives at each number of individual and cell combination. In this sense, it is true that pseudobulk approaches have mis-calibrated confidence intervals, obtaining fewer false positives than expected at a 0.05 p-value cut-off. The potential issue with this is, that given pseudobulk methods’ conservative 95% confidence intervals, it would be expected to have a higher type 2 error than other methods. However, our ROC analysis disproves this. It shows how, at equal type 1 error rates, pseudobulk mean has the lowest type 2 error rate of all tested methods (Figure 1, Supplementary Figure 2).

All tests to this point have been on simulated with an equal number of cells in each sample. However, in real datasets there would never be the case. To mirror this, we simulated data with an imbalanced number of cells between case and controls. Pseudobulk mean outperformed all other approaches on this analysis (Supplementary Figure 3). The pseudobulk approach which aggregated by averaging rather than taking the sum appears to be the top performing overall. However, it is worth noting that hierarchicell does not normalise the simulated datasets before passing to the pseudobulk approaches. This is a standard step in such analysis to account for differences in sequencing depth and library sizes^5^. This approach was taken by Zimmerman *et al*. as their data is simulated one independent gene at a time without considering differences in library size. The effect of this step is more apparent on the imbalanced number of cells where pseudobulk sum’s performance degraded dramatically. Pseudobulk mean appears invariant to this missing normalisation step because of averaging’s own normalisation effect. Importantly, this was a flaw in the simulation software strategy and does not show an improved performance of pseudobulk mean over sum. We believe this approach also effected pseudobulk sum’s performance on the different proportions of differentially expressed genes (Figure 1).

Pseudobulk approaches were also found to be the top performing approaches in a recent review by Squair *et al*.,^4^. Notably, the pseudobulk method used here; DESeq2^5^, performed worse than other pseudobulk models in Squair *et al*.,’s analysis and so their adoption may further increase the performance of pseudobulk approaches o n our dataset. Conversely, Squair *et al*., did not consider all models included in our analysis or the different forms of pseudobulk aggregation. Therefore, our results on sum and mean pseudobulk extend their findings and indicate that mean aggregation may be the best performing. However, the reader should be cognisant that the lack of a normalisation step based on the flaw in the simulation software strategy likely causes the increased performance of mean over sum aggregation. Further, the use of simulated datasets in our analysis may not accurately reflect the differences between individuals that are present in biological datasets. Thus, despite both our results and those reported by Squair *et al*., there is still room for further analysis, benchmarking more models, including different combinations of pseudobulk aggregation methods and models, on more representative simulated datasets and biological datasets to identify the optimal approach. Specifically, we would expect pseudobulk sum with a normalisation step to outperform pseudobulk mean since it can account for the intra-individual variance which is otherwise lost with pseudobulk mean but this should be tested, including on imbalanced datasets and at consistent type 1 error rates.

## Conclusion

In conclusion, our results demonstrate that pseudobulk approaches are the best performing for the analysis of single-cell expression data based on power at equivalent type 1 error rates and on MCC for both balanced and imbalanced number of cells, from this simulated dataset.

## Supplementary Figures and Tables

**Supplementary Figure 1:**
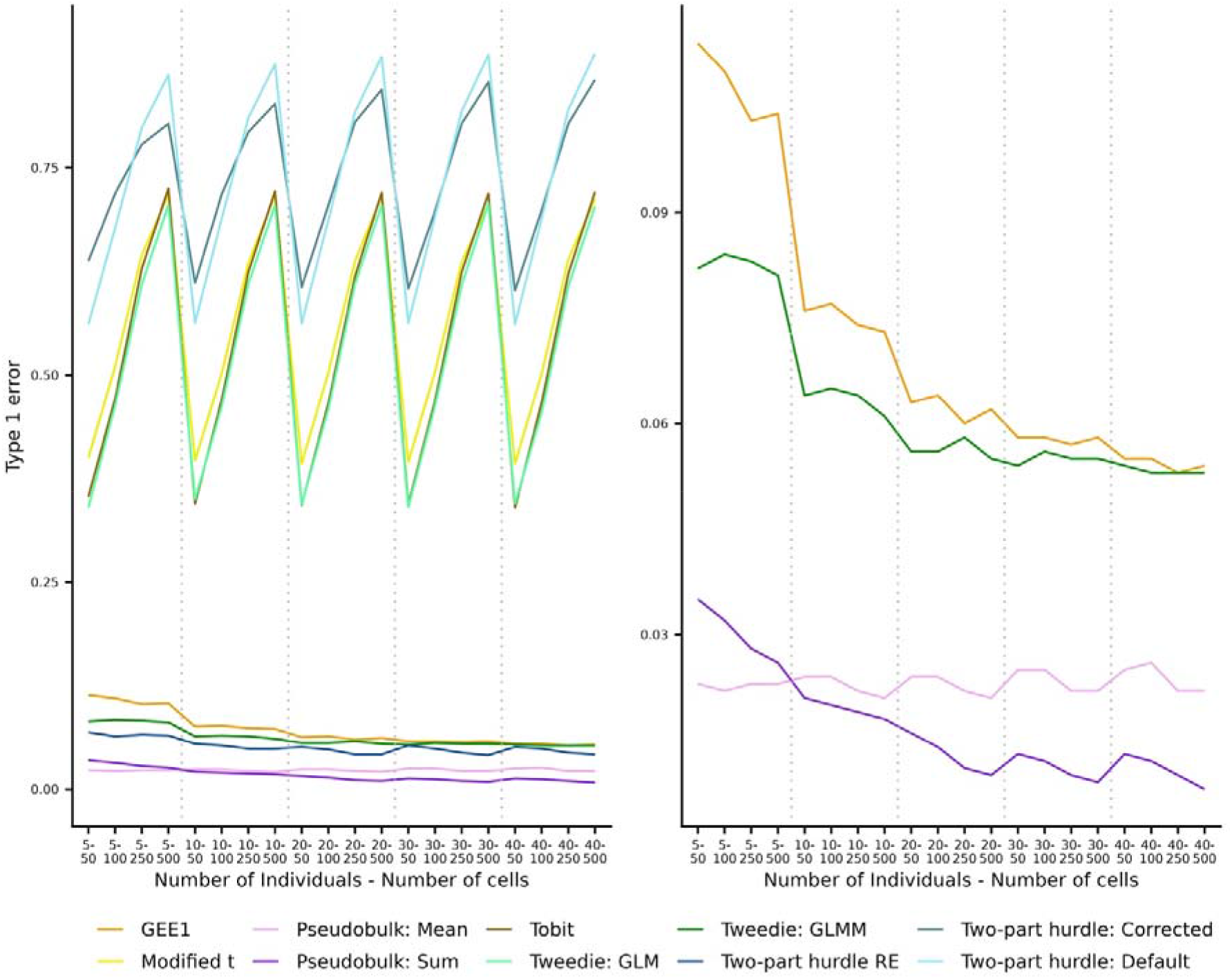
The average type 1 error from the 20,000 iterations; 50 runs for each of the 5 to 40 individuals and 50 to 500 cells at a p-value cut-off of 0.05 on 5,000 genes reported by Zimmerman et al.^1^. Left shows all benchmarked models whereas right focuses on the top four approaches. The different models are pseudoreplication approaches; ‘Modified t’, ‘Tobit’, ‘Two-part hurdle: Default’, ‘Two-part hurdle: Corrected’, ‘GEE1’, ‘Tweedie: GLM’, pseudobulk approaches; ‘Pseudobulk: Mean’, ‘Pseudobulk: Sum’ and mixed model approaches; ‘Tweedie: GLMM’ and ‘Two-part hurdle: RE’.

**Supplementary Figure 2:**
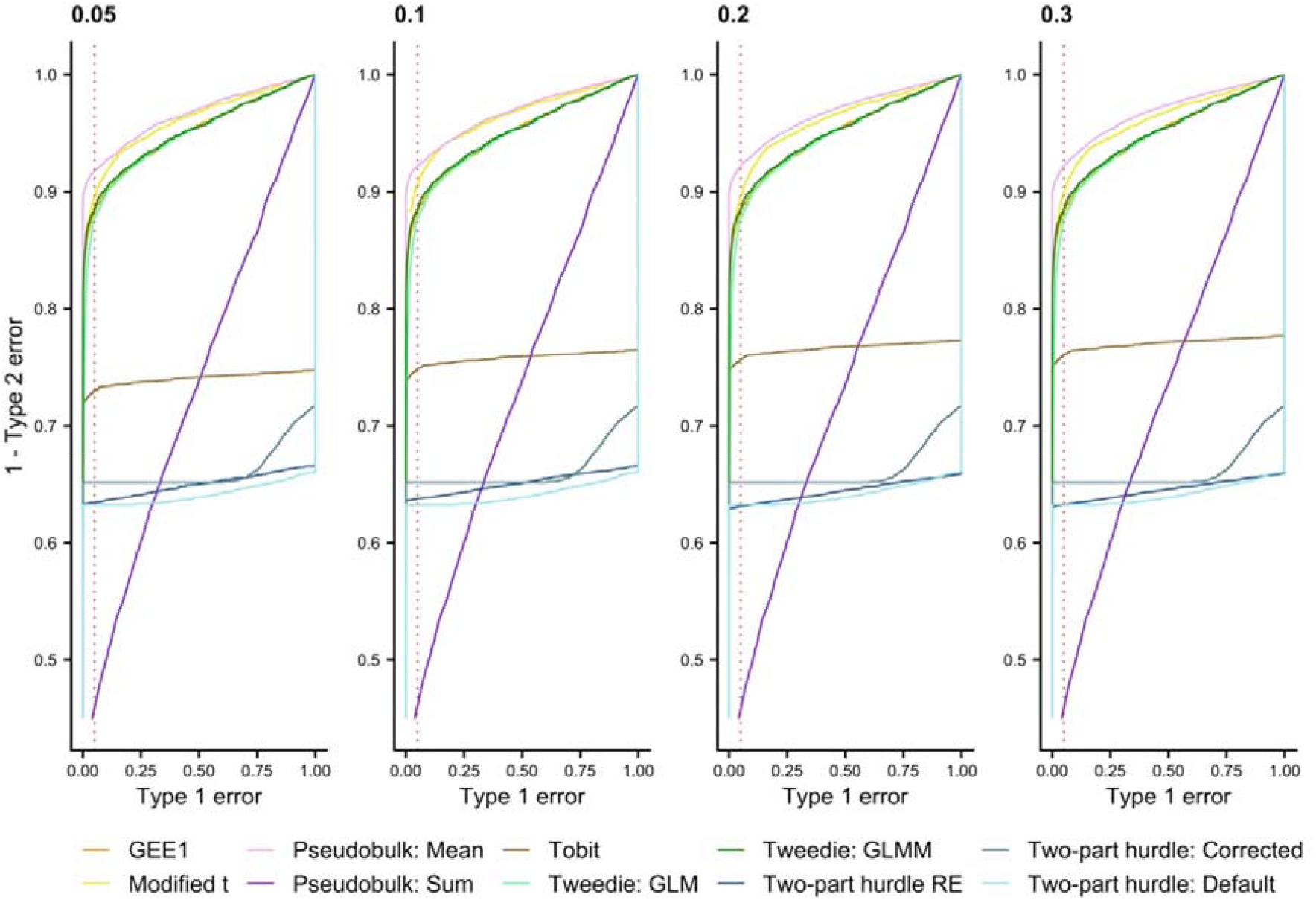
Pseudobulk mean is best performing at a constant type 1 error rate. The four images give the receiver operating chracteristics (ROC) curve across 50 runs each for different proportions of simulated differentially expressed genes (DEGs) - 0.05, 0.1,0.2,0.3. 20 individuals were simulated for case and controls, each with 100 cells. The proportion of DEGs are of the 5,000 non-DEGs simulated. The sensitivity (1 – type 2 error) of the different approaches at a 0.05 type 1 error value are highlighted by the red dashed line. The different models are pseudoreplication approaches; ‘Modified t’, ‘Tobit’, ‘Two-part hurdle: Default’, ‘Two-part hurdle: Corrected’, ‘GEE1’, ‘Tweedie: GLM’, pseudobulk approaches; ‘Pseudobulk: Mean’, ‘Pseudobulk: Sum’ and mixed model approaches; ‘Tweedie: GLMM’ and ‘Two-part hurdle: RE’.

**Supplementary Figure 3:**
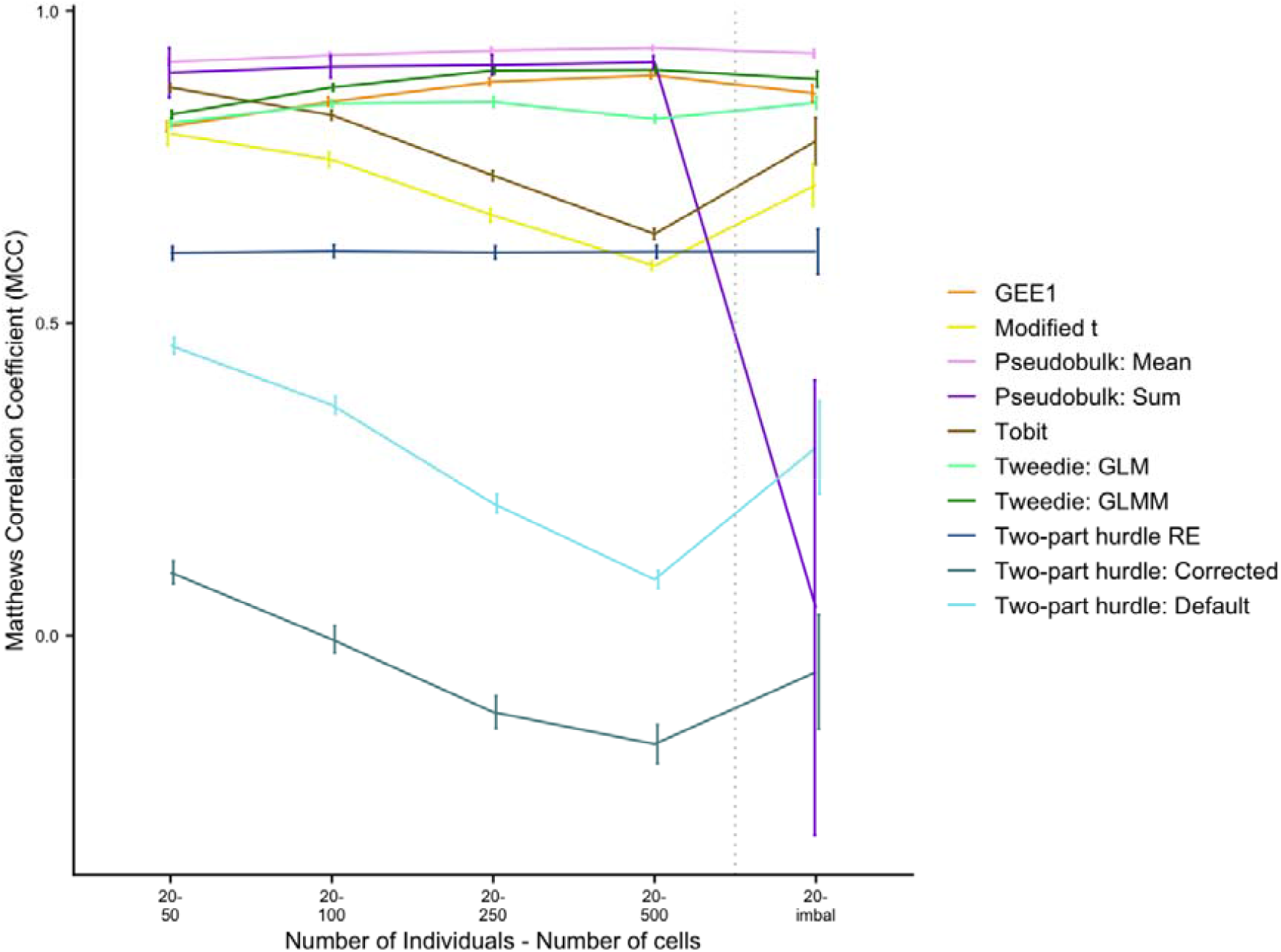
The average Matthews correlation coefficient of all benchmarked models across all balanced number of cells and the imbalanced number of cells for 20 individuals; 50 runs for each at a p-value cut-off of 0.05 on 5,000 genes. The number of cells were randomly chosen using a gamma distribution with shape 4 and scale 45 separately for cases and controls to produce the imbalanced dataset (giving a mean 150-200 cells). The error bars give 1 standard deviation around the mean. The different models are pseudoreplication approaches; ‘Modified t’, ‘Tobit’, ‘Two-part hurdle: Default’, ‘Two-part hurdle: Corrected’, ‘GEE1’, ‘Tweedie: GLM’, pseudobulk approaches; ‘Pseudobulk: Mean’, ‘Pseudobulk: Sum’ and mixed model approaches; ‘Tweedie: GLMM’ and ‘Two-part hurdle: RE’.

**Supplementary Table 1:**
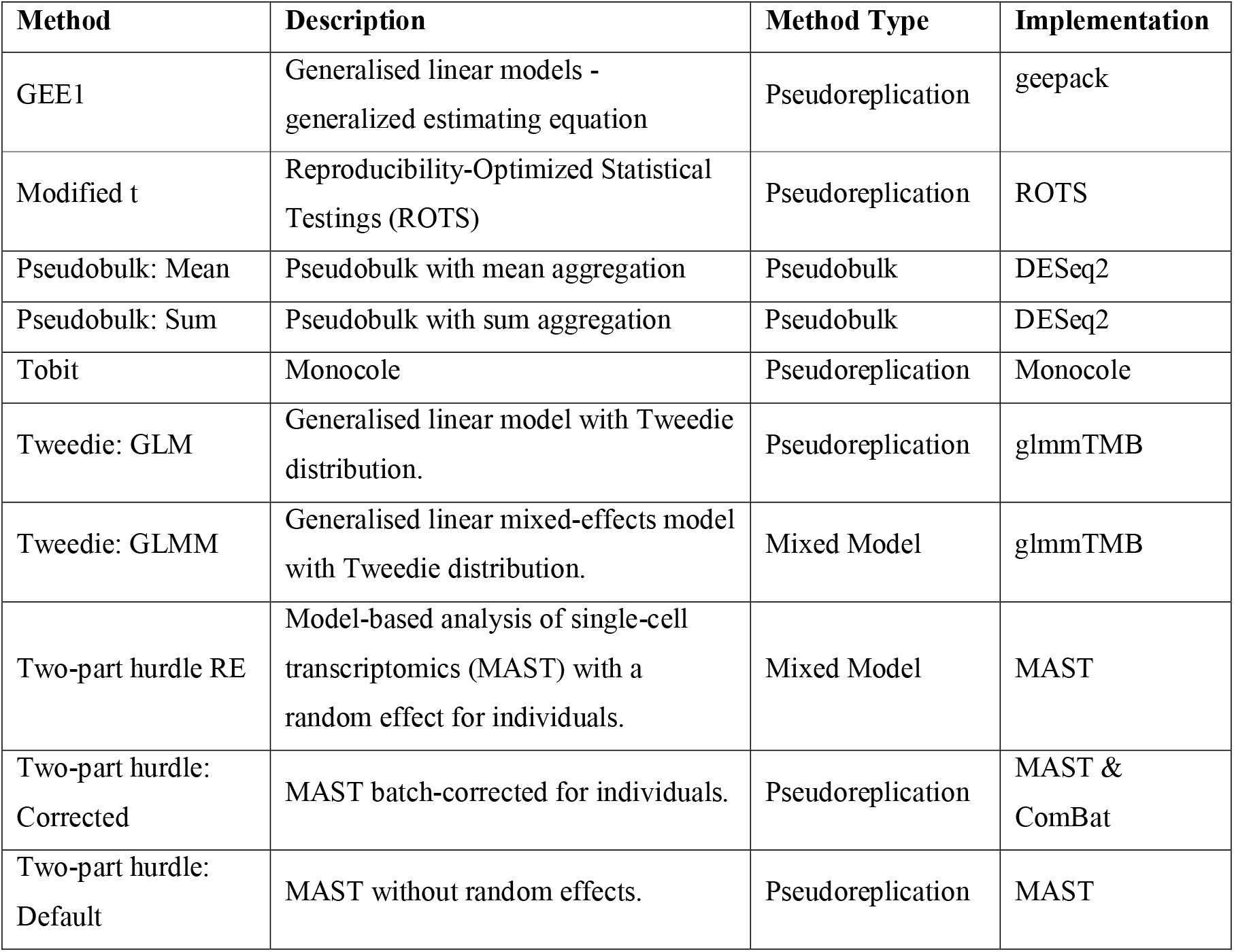
The different methods benchmarked in the analysis with their implementation approach and methodological types.

| Method | Description | Method Type | Implementation |
| --- | --- | --- | --- |
| GEE1 | Generalised linear models - generalized estimating equation | Pseudoreplication | geepack |
| Modified t | Reproducibility-Optimized Statistical Testings (ROTS) | Pseudoreplication | ROTS |
| Pseudobulk: Mean | Pseudobulk with mean aggregation | Pseudobulk | DESeq2 |
| Pseudobulk: Sum | Pseudobulk with sum aggregation | Pseudobulk | DESeq2 |
| Tobit | Monocle | Pseudoreplication | Monocle |
| Tweedie: GLM | Generalised linear model with Tweedie distribution. | Pseudoreplication | glmmTMB |
| Tweedie: GLMM | Generalised linear mixed-effects model with Tweedie distribution. | Mixed Model | glmmTMB |
| Two-part hurdle RE | Model-based analysis of single-cell transcriptomics (MAST) with a random effect for individuals. | Mixed Model | MAST |
| Two-part hurdle: Corrected | MAST batch-corrected for individuals. | Pseudoreplication | MAST & ComBat |
| Two-part hurdle: Default | MAST without random effects. | Pseudoreplication | MAST |

**Supplementary Table 2:**
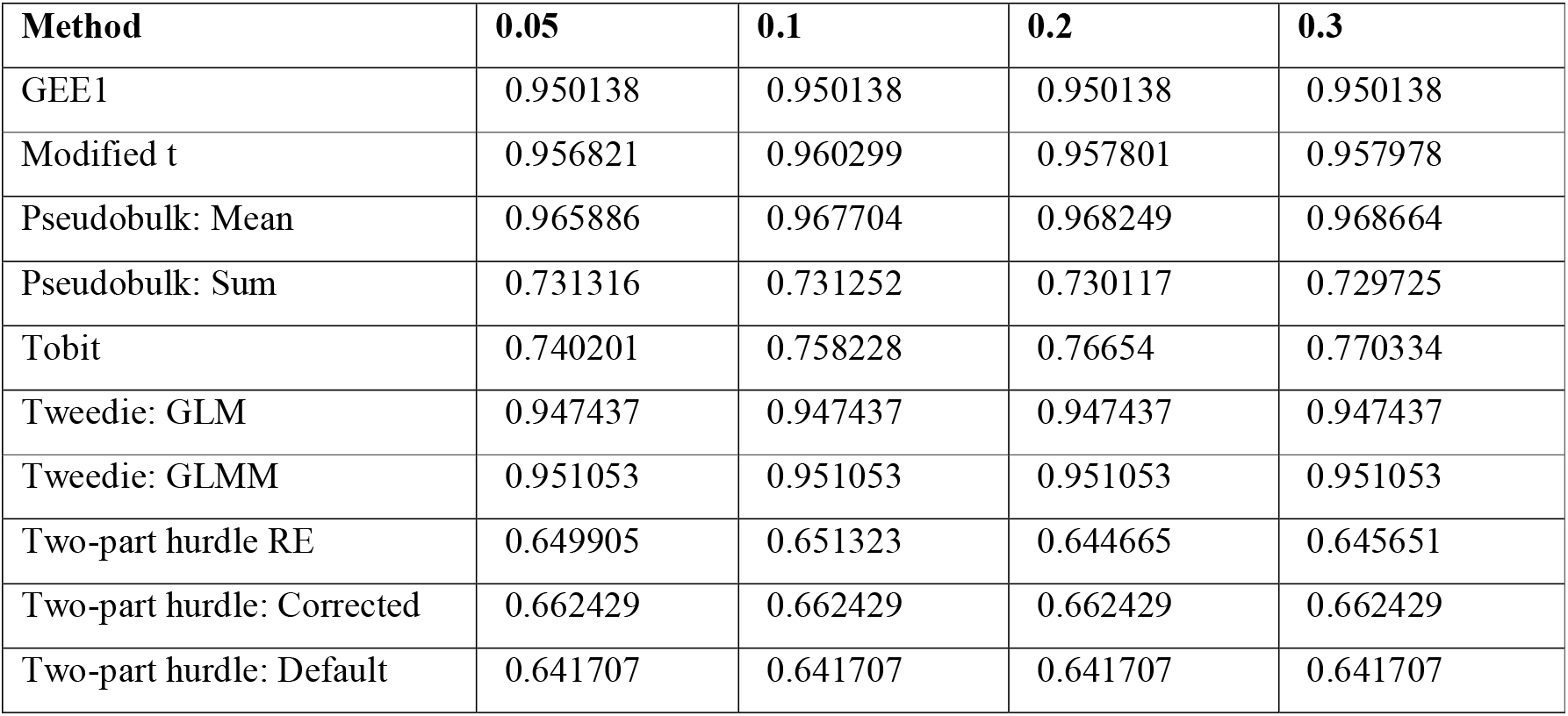
The area under the receiver operating characteristics curve (AUC) of all benchmarked models across different proportions of simulated differentially expressed genes (DEGs) - 0.05, 0.1,0.2,0.3. 20 individuals were simulated for case and controls, each with 100 cells. The proportion of DEGs are of the 5,000 non-DEGs simulated. The values are rounded to six decimal places. The different models are pseudoreplication approaches; ‘Modified t’, ‘Tobit’, ‘Two-part hurdle: Default’, ‘Two-part hurdle: Corrected’, ‘GEE1’, ‘Tweedie: GLM’, pseudobulk approaches; ‘Pseudobulk: Mean’, ‘Pseudobulk: Sum’ and mixed model approaches; ‘Tweedie: GLMM’ and ‘Two-part hurdle: RE’.

